# Polyommatine blue butterflies reveal unexpected integrity of the W sex chromosome amid extensive chromosome fragmentation

**DOI:** 10.1101/2024.06.25.600692

**Authors:** Monika Hospodářská, Anna Chung Voleníková, Petr Koutecký, Roger Vila, Gerard Talavera, Irena Provazníková, Martina Dalíková, Petr Nguyen

**Affiliations:** University of South Bohemia, Faculty of Science, České Budějovice 370 05, Czech Republic; Biology Centre of the Czech Academy of Sciences, Institute of Entomology, České Budějovice, 370 05, Czech Republic; Institut de Biologia Evolutiva, Consejo Superior de Investigaciones Científicas and Universitat Pompeu Fabra, Barcelona, Spain; Institut Botànic de Barcelona (IBB), Consejo Superior de Investigaciones Científicas and Consorci Museu de Ciències Naturals de Barcelona, Barcelona, Spain; European Molecular Biology Laboratory, Heidelberg, Germany; University of Kansas, Lawrence, Kansas, USA

## Abstract

Chromosomal rearrangements are crucial in speciation, acting as barriers to gene flow. Holocentric chromosomes, such as those in Lepidoptera, can facilitate karyotype changes. Despite chromosome fusions being more common, speciation events are mostly linked to fissions. Notable karyotypic variation is observed in three clades of the subfamily Polyommatinae (Lycaenidae), with chromosome numbers ranging from n = 10 to n = 225. This study used flow cytometry and molecular cytogenetic analyses to investigate genome sizes and karyotypes in several species of the genera *Polyommatus* and *Lysandra* with derived and modal chromosome numbers. The findings show no support for polyploidy, supporting karyotypic diversification via fragmentation of chromosomes. Species with high chromosome numbers have larger genomes, which indicates a potential role of mobile elements but contradicts the hypothesis of holocentric drive. Telomeric signals were detected at the ends of fragmented chromosomes. No interstitial telomeric sequences were detected on autosomes. Interstitial telomeric signals on sex chromosomes, however, revealed multiple sex chromosome systems in *Polyommatus dorylas* and *Polyommatus icarus*, with two karyotype races differing in sex chromosome constitution in the latter. Pool-seq and coverage analyses indicated shared fusion of sex chromosomes with an autosome bearing the rDNA locus, followed by a fusion with chromosome 20 in the Czech population. Notably, the W chromosome resists fragmentation, likely due to epigenetic silencing protecting it from activity of mobile elements.

## Introduction

Chromosomal rearrangements supposedly play a crucial role in speciation as they can serve as barriers to gene flow (Faria and Navarro 2010). Theoretical models of chromosomal speciation have traditionally concerned monocentric chromosomes, which have a localized centromeric region. However, holocentric chromosomes, which lack a distinct centromere, may facilitate karyotype change, and thus potentially impact the process of speciation (Lucek et al. 2022). Holocentric chromosomes have evolved multiple times and represent a significant portion of biodiversity (Drinnenberg et al. 2014, Escudero et al. 2016).

Moth and butterflies (Lepidoptera) comprising more than 160 000 species (van Nieukerken et al. 2011) represent the most species rich holocentric lineage. Only ca. 1.5% of species were karyotyped (de Vos et al. 2020) and few were cytogenetically analyzed in detail due to uniformity of their small metaphase chromosomes which complicated identification of individual chromosomes (de Prins and Saitoh 2003). Recent advances in sequencing technologies provided robust phylogenetic framework and brought genomes to chromosome level, thus offering an unprecedented opportunity to better understand the role of chromosomal changes in lepidopteran speciation (Augustijnen et al. 2024, Wright et al. 2024).

Lepidopteran genome organization including synteny of genes is highly conserved across distant lineages (de Vos et al. 2020, Wright et al. 2024). The most frequent chromosome number n = 31 is also ancestral (Traut et al. 2023, Van’T Hof et al. 2013, Wright et al. 2024) and extensive karyotype changes are limited only to few groups (de Vos et al. 2020, Wright et al. 2024). Phylogenetic analyses of diversification indicated a positive association between speciation rates and karyotype evolution in Lepidoptera (Talavera et al. 2013a, de Vos et al. 2020, Augustijnen et al. 2024). Although chromosome fusions are more common than fissions, cladogenetic chromosomal changes associated with speciation events mostly correspond to fissions (de Vos et al. 2020, Augustijnen et al. 2024).

Blue butterflies of the family Lycaenidae possess the greatest interspecific karyotype variation with chromosome numbers ranging between n = 10–225 (Kandul et al. 2004) and no evidence of polyploidization (Kandul et al. 2007, Wright et al. 2024). Their modal chromosome number was first reduced to n = 24 in the lycaenid last common ancestor (Robinson 1971, Wright et al. 2024). The majority of blue butterflies with derived chromosome numbers belong to three closely related lineages within the subtribe Polyommatina, genus *Lysandra* (n = 24–92) and previously recognized subgenera *Agrodiaetus* (n = 10–125) and *Plebicula* (n = 134–225), which are now classified under the genus *Polyommatus* (Kandul et al. 2004, Talavera et al. 2013a). Intraspecific variation in chromosome number has been reported as well. For example, n = 87–88 was observed in Spanish, French and Italian populations of *Lysandra coridon* (Schmitt and Seitz 2001), while specimens from the Balkans possess n = 90–92 chromosomes. Kandul et al. (2004) hypothesized that karyotypic diversification in the subgenus *Agrodiaetus* (n = 10–125) was due to activity of mobile elements horizontally transferred between the closely related species. Comparative phylogenetic analysis in the subgenus *Agrodiaetus* showed that chromosomal numbers are more diverse between young sympatric species than in corresponding allopatric pairs. This implies that chromosomal rearrangements play direct role in speciation process (Kandul et al. 2007). Increased diversification rate was confirmed in *Lysandra* spp. (Kandul et al. 2007, Talavera et al. 2013b). However, analysis of Vershinina and Lukhtanov (2017) revealed a correlation between the amount of karyotype differences and length of phylogenetic branches.

Interestingly, two large chromosomes were observed in species with high chromosome numbers such as *Polyommatus* (*Agrodiaetus*) *damone damone* or *P. atlantica* (Lukhtanov 2015, Lukhtanov et al. 1997). According to Ennis (1976) the large chromosomes result from autosomal fusions, while White (1946) supposed that the large chromosomes observed in blue butterflies with high chromosome number correspond to sex chromosomes. Indeed, comparative genomics show that the sex chromosome Z remains intact in lineages with extensive chromosome fragmentation (Wright et al. 2024). Moreover, recent studies have shown that unlike in female heterogametic vertebrates (Pennell et al. 2015), sex chromosome-autosome fusions are quite common in Lepidoptera including representatives of the family Lycaenidae (Carabajal Paladino et al. 2019, Pazhenkova and Lukhtanov 2023; Wright et al. 2024) and these sex chromosome turnovers may be driven by sexual antagonistic selection (Mora et al. 2024). Yet, sex chromosomes remain generally understudied in female heterogametic taxa (Ellegren 2011, Tomaszkiewicz et al. 2017) and particularly female limited W chromosomes are not well represented in lepidopteran chromosome-level assemblies due to their high repeat content and lower sequencing depth compared to autosomes (Carey et al. 2022, Wright et al. 2024).

Here we used flow cytometry in representatives of the genera *Polyommatus* and *Lysandra* (Fig. 1) which differ in their chromosomal numbers, namely, *P. icarus*, *P. escheri* and *P. thersites* with modal chromosome number (n = 23/24; Nguyen et al. 2010, Kandul et al. 2004, Robinson 1971) and representatives of genera with high chromosomes numbers *L. bellargus* (n = 45; Nguyen et al. 2010), *L. coridon* (n = 87–97; (Schmitt and Seitz 2001), *L. hispana* (n = 84; Descimon and Mallet 2010), *P.* (*Agrodiaetus*) *ripartii* (n = 90; Munguira et al. 1995), *P.* (*A.*) *fulgens* (n = 103; Munguira et al. 1995) and *P.* (*Plebicula*) *dorylas* (n = 149– 151; (Wiemers 2003). We determined their genome sizes to directly test whether polyploidization or repeat expansion played a role in their karyotype evolution. We further performed molecular cytogenetic analyses of *P. icarus, P. escheri*, *L. bellargus, L. coridon* and *P. dorylas* with particular focus on sex chromosomes. Our results show little variation in repeat content and counterintuitive conservation of sex chromosomes with the W chromosome being the most stable element of lycaenid karyotypes.

**Figure 1.**
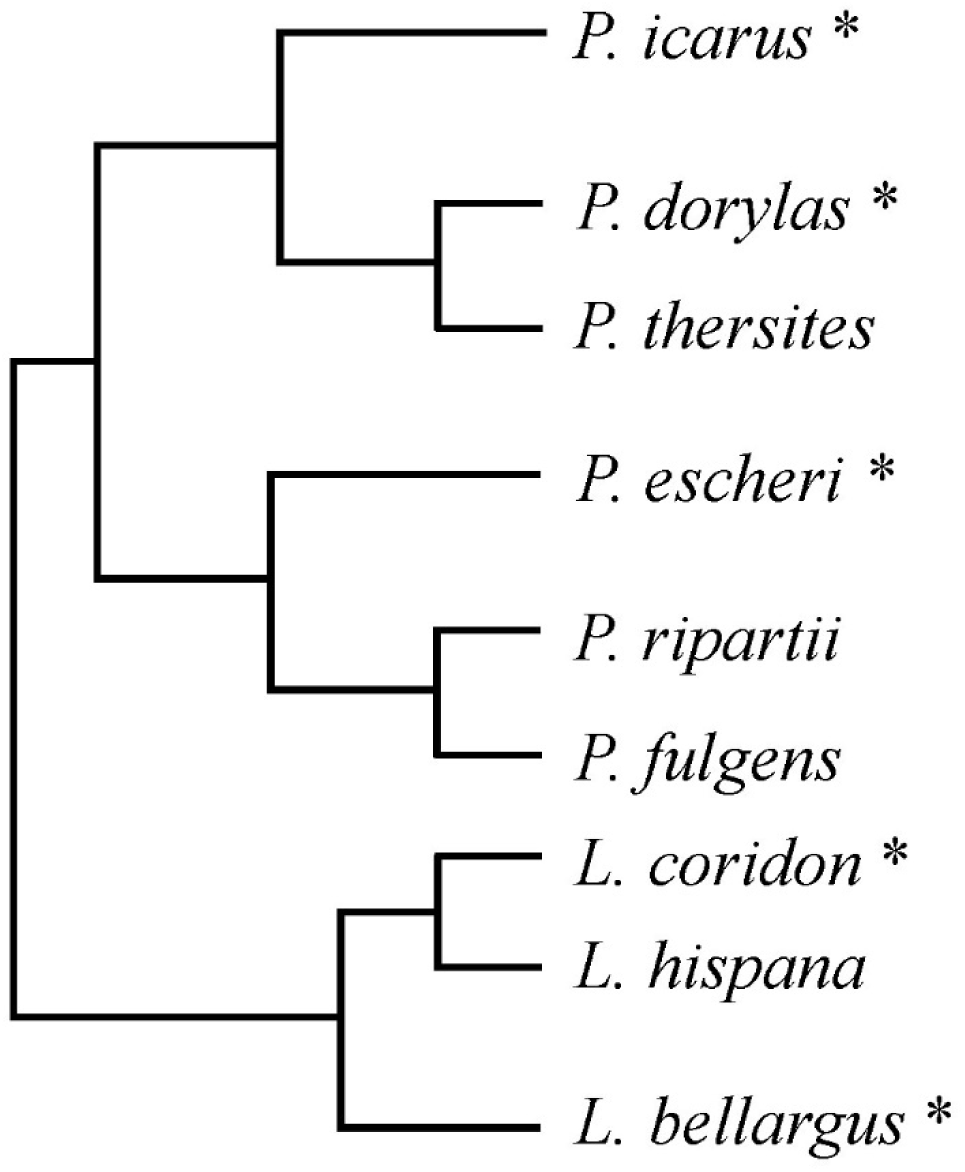
Phylogenetic relationships of *Polyommatus* and *Lysandra* species under study (according to Kandul et al. 2004, Wiemers et al. 2010, Talavera et al. 2013b). * Species used in molecular cytogenetic analyses.

## Materials and Methods

### Sample collecting

For hybridization experiments we sampled 5 representatives of the subfamily Polyommatinae. Adult females of *Polyommatus icarus*, *Lysandra bellargus* and *Lysandra coridon* were collected in the Czech Republic, namely in the surroundings of České Budějovice, in Hády stone quarry in Brno and around Týnčany village in Central Bohemian Region, respectively. Larvae of *Polyommatus dorylas* species were obtained from Pavel Skála from maintained breeding which originates from wide stock from Czech Republic. The last species, *Polyommatus escheri*, was collected in the Montseny area near Barcelona, Spain. Hatched larvae from fertilized females were reared on corresponding host plants, *Lotus corniculatus* for *P*. *icarus*, *Coronilla varia* for *L*. *bellargus* and *L*. *coridon*, *Anthyllis vulneraria* for *P*. *dorylas* and *Astragalus monspessulanus* for *P*. *escheri*, at room temperature and normal day/night regime. Gonads and imaginal discs of 5^th^ instar larvae were used to make chromosomal preparations and the remaining tissue was frozen in liquid nitrogen and stored at −80 °C until DNA extraction.

Estimation of genome size was done on adult males collected in different areas. Males of *P*. *icarus* were collected in Suchdol and Lužnicí in South Bohemia, Czech Republic, and moreover in the Montseny area near Barcelona, Spain, along with *P*. *escheri*, *P*. *thersites*, *P*. *ripartii* and *P*. *fulgens*. *P. dorylas* and P. *hispana* were collected in Coll de Pal and the Figols municipality area, Spain, respectively. The last species *L*. *coridon* was collected in the Hády stone quarry in Brno, Czech Republic. All individuals were frozen in liquid nitrogen and stored at −80 °C until further use. As standard we used the WT-C laboratory wild-type strain of *Ephestia kuehniella* (Marec 1990). Larvae of this species were kept on wheat meal with brewer’s yeast in a room with a temperature of 21 ± 1 °C, at a 12/12 h (light/dark) regime.

### Genome size estimation

We estimated the size of selected blue butterfly’s genome using flow cytometry according to Buntrock et al. (2012) with some modifications. Briefly, the head (or 1/2) of a single tested specimen and the head of a single *E*. *kuehniella* standard (1C = 0.45 pg; Buntrock et al. 2012) were homogenized together in 1 ml of Tris-MgCl_2_ buffer (consisting of: 100 Mm Tris-HCl pH 7.5, 2 mM MgCl_2_ and 1% Triton X-100) with razor blade and filtered through a 42 µm filter (Uhelon 130 T). Cells were then stained by propidium iodide of final concentration 50 µg/ml for at least 15 min. Three male samples of each species were prepared independently. Measurements were done in the CyFlow SL (Partec, Münster, Germany) flow cytometer equipped with a green diode-pumped solid-state laser emitting an exciting light at 532 nm and the fluorescence of 5 000 particles was recorded. Evaluation of histograms was done with FlowJo software, version 10 (FlowJo, LLC, Ashland, OR, USA). The amount of DNA was determined as the ratio of the 2C mean of the sample with the 2C mean for standard *E*. *kuehniella* multiplied by genome size of the standard.

### Molecular-cytogenetics analyses

Mitotic and meiotic chromosome spreads were made from imaginal discs and ovaries of last instar larvae following Mediouni et al. (2004) with slight modifications listed in Šíchová et al. (2013). The preparations were dehydrated in an ethanol row (70-80-100 %, 30–60 s each) and along with the rest of larvae body, which was frozen with liquid nitrogen, stored at -80 °C.

The frozen bodies were used for DNA extraction with hexadecyltrimethylammonium bromide (CTAB) according to Winnepenninckx et al. (1993). Extracted male genomic (gDNA) was amplified according to manufactureŕs protocol by GenomiPhi HY kit (Ge Healthcare). The concentration of DNA amount was measured with the Qubit 3.0 fluorometer (Thermo Fisher Scientific, Waltham, MA, USA).

Telomeric template DNA (TTAGG)_n_ was generated using non-template polymerase chain reaction (PCR) with two oligomers 5′-TAGGTTAGGTTAGGTTAGGT-3′ and 5′-CTAACCTAACCTAACCTAAC-3′ (Sahara et al. 1999). Reactions were caried out in 100 µl reaction volumes containing 0.5 µM each of primers, 1x Ex Taq pufr, 0.8 mM dNTP mix and 2 U Ex Taq polymerase (TaKaRa, Otsu, Japan). The thermal profile of the reaction started with initial stage 90 s at 94 °C which was followed by 30 cycles of 45 s denaturation at 94 °C, 30 s annealing at 52 °C and 60 s synthesis at 72 °C. Final synthesis took 10 min at 72 °C. PCR product was precipitated (30 min at -80 °C) by adding 1/10 of volume of sodium acetate and 2.5 times volume of 100% ethanol.

Fragment of the gene for 18S ribosomal RNA was amplified by PCR from DNA of codling moth, *Cydia pomonella*, with specific primers 5′-CGATACCGCGAATGGCT-CAATA-3′ and 5′-ACAAAGGGCAGGGACGTAATCAAC-3′ according to Fuková et al. (2005) with some modifications. Reactions were caried out in 25 µl reactions volumes containing 10 ng template DNA, 0.5 µM of each primer, 0.2 mM dNTP mix, 1x Ex Taq buffer and 2 U Ex Taq polymerase (TaKaRa, Otsu, Japan). The thermal profile started with initial stage 94 °C for 3 min which was followed by 30 cycles of 1 min denaturation at 94 °C, 1 min annealing at 52 °C, 1 min synthesis at 72 °C. The final synthesis took 5 min at 72 °C. Both obtained telomeric DNA and 18S ribosomal DNA (rDNA) were then purified by Wizard SV Gel and PCR Clean-Up System (Promega, Madison, WI, USA) and quantified by Qubit 3.0 fluorometer.

Hybridization probes were prepared from female genomic DNA (gDNA), (TTAGG)_n_ telomeric DNA and 18S rDNA, which were labelled by fluorochrome Cy3-dUTP (Jenna Bioscience, Jena, Germany), Fluorescein-12-dUTP (Jenna Bioscience, Jena, Germany) or biotin-16-dUTP (Hoffmann-La Roche, Basilej, Switzerland) using the nick translation method. The 40 µl reaction contained 2 µg of DNA, 1x nick translation buffer (5 mM Tris-HCl, 0.5 mM MgCl_2_, 0.0005% BSA; pH 7.5), 10 mM β-mercaptoethanol, 50 µM dATP, 50 µM dCTP, 50 µM dGTP, 10 µM dTTP, 20 µM Cy3-dUTP/Fluorescein-12-dUTP/biotin-16- dUTP, 40 U DNA polymerase I (Thermo Fisher Scientific, Waltham, MA, USA) and 0.01 U DNase I (RNase-free, Thermo Fisher Scientific).

The incubation at 15 °C varies according to input DNA with gDNA being labelled for 3.5 h and (TTAGG)_n_ telomeric DNA and rDNA for 1 h. Finally, the reaction was stopped by adding 1x loading buffer (50% glycerol, 250 mM EDTA, 5.9 mM bromphenol blue).

Some of chromosomal preparations were stained by 2.5% lactic acetic orcein (LAO, 2.5% orcein in 30% acetic acid and 30% milk acid) to detect the W-specific heterochromatin. After 55 min incubation at room temperature, lactic acetic orcein was washed away by distilled water. Preparations were inspected with phase microscope Zeiss Axioplan 2 (Carl Zeiss, Jena, Germany) with cooled CDC camera XM10. After that the immersion oil was washed away by incubation 1 min in xylen and 1 min in benzene which was washed by distilled water. The incorporated lactic acetic orcein was washed away by methanol:acetic acid (3:1) so the preparations could be used for hybridization experiments as well.

For detection of the W sex chromosome, we used genomic *in situ* hybridization (GISH) with female whole genome labelled probe. The experiment was done according to Sahara et al. (1999) with some modifications. Firstly, preparations were treated by RNase A (100 µg/ml) for 1 h at 37 °C and washed twice for 15 min in 2x SSC. We used 5x Denhardt’s solution (0.1% Ficoll, 0.1% polyvinylpyrrolidone, 0.1% bovine serum albumin) as blocking solution in which preparations were incubated for 30 min in water bath at 37 °C. Chromosomes were denatured by 70% formamide in 2x SSC for 3.5 min at 68 °C and then immediately dehydrated in cold ethanol row. For each slide we prepared 10 µl of hybridization mix which contained 250 ng labelled female gDNA, 3 µg unlabelled competitor DNA, 25 µg sonicated salmon sperm DNA (Sigma-Aldrich, Munich, Germany), 50% formamide and 10% dextran sulfate in 2x SSC. As the competitor DNA we used male gDNA, which was first fragmented by incubation 20 min at 99 °C. The slides were hybridized for 3 days at 37 °C. Posthybridization washes were done in 1% Triton X-100 in 0.1x SSC for 5 min at 62 °C and 2 min in Ilfotol (Ilford) at room temperature. The preparations were counterstained with 0.5 mg/ml DAPI and mounted in antifade based on DABCO.

The same chromosomal preparations were also used for *in situ* hybridization with (TTAGG)_n_ telomeric probe according to reprobing protocol described in (Nguyen et al. 2013). Cover slips were removed, and preparations were washed two times in 2x SSC for 5 min. Chromosomal preparations were postfixed by 10 min incubation in 4% paraformaldehyde in 2x SSC at room temperature and then washed three times in 2x SSC. Simultaneously the genomic probe was washed, and chromosomes were denatured by incubation in 50% formamide in 1% Triton-X100 in 0.1x SSC for 10 min at 70 °C. The slides were immediately dehydrated with the ethanol series. These preparations were ready for new 10 µl hybridization mix which consisted of 150 ng labelled (TTAGG)_n_ telomeric probe, 25 μg sonicated salmon sperm DNA (Sigma-Aldrich, Munich, Germany), 50% formamide and 10% dextran sulfate in 2x SSC.

Alternatively, chromosomal preparations were hybridized with the 18S rDNA probe. We used 10 µl of hybridization mix per one slide which contained 30 ng biotin-labelled 18S rDNA, 25 μg sonicated salmon sperm DNA (Sigma-Aldrich, Munich, Germany), 50% formamide and 10% dextran sulfate in 2x SSC. The signal was detected with Cy3-conjugated streptavidin (Jackson ImmunoResearch Laboratories Inc, West Grove, PA, US) and enhanced with biotinylated anti-streptavidin (Vector Labs. Inc, Burlingame, CA, USA) again detected with Cy3-conjugated streptavidin as in Fuková et al. (2005).

All hybridization experiments were documented with fluorescent microscope Zeiss Axioplan 2 (Carl Zeiss, Jena, Germany) with cooled CDC camera XM10. Digital images of each separate fluorescent dye were collected using cellSens 1.9 software (Olympus Europa Holding, Hamburg, SRN) and subsequently pseudocoloured and superimposed with Adobe Photoshop, version 9.0.

### Pool-seq data analysis

*Polyommatus icarus* males and females were collected across South Bohemia. Offspring of the collected females was probed by GISH to confirm their WZ_1_Z_2_ sex chromosome constitution. In total, gDNA from 30 males and 30 females with WZ_1_Z_2_ were mixed into sex-specific pools, which were used for preparation of Illumina libraries and sequenced. The raw reads were quality checked with FastQC v0.11.5 (Andrews et al. 2010). Filtering and trimming were done with Trimmomatic v0.36 (Bolger et al. 2014) with the following parameters: “LEADING:3 TRAILING:3 SLIDINGWINDOW:4:25 MINLEN:100”. The female and male pools were mapped separately to the genome using BWA-MEM v0.7.17 (Li and Durbin 2009) with the default parameters. As a reference genome, we used *Polyommatus icarus* chromosome level genome assembly GCA_937595015.1 (Lohse and Vila 2023). Resulting bam files were sorted by coordinate (“SortSam SORT_ORDER=coordinate”) and the PCR duplicates were removed (“MarkDuplicates REMOVE_DUPLICATES=true REMOVE_SEQUENCING_DUPLICATES=true”) with Picard Toolkit v2.22.1 (https://broadinstitute.github.io/picard/). A file with the nucleotide composition of all genomic positions was generated using the pileup function of the software Pooled Sequencing Analysis for Sex Signal v3.1.0 (PSASS; DOI: zenodo.org/records/4442702). The PSASS was also used to identify non-overlapping 50 kb windows enriched in sex-specific SNPs, using analyse function with the following parameters: “--min-depth 10 --freq-het 0.5 --range-het 0.15 --freq-hom 1 --range-hom 0.05 --window-size 50000 --output-resolution 50000 --group-snps”.

### Coverage analysis

In total, three females and five males, which were used also in Pool-seq analysis, were sequenced as short reads by Illumina. The quality was checked with FastQC v0.11.5 (Andrews et al. 2010). Poly-G artifacts of two-channel sequencing Illumina system were trimmed by Cutadapt v0.1.15 (Martin 2011) using “--nextseq-trim=20 --minimum-length=100” options. Then, we used Trimmomatic v0.36 (Bolger et al. 2014) for quality filtering and trimming with following parameters: “SLIDINGWINDOW:4:20 MINLEN:75 HEADCROP:4 CROP:140”. Repetitive sequences were identified in reference genome assembly by RepeatModeler v1.0.11 (Smit and Hubley 1996-2010) and annotated with the NCBI search by RepeatMasker v4.0.7 (Smit et al. 2008-2015; both available at http://www.repeatmasker.org). Trimmed reads were mapped to the masked reference assembly via Bowtie2 v2.2.9 (Langmead and Salzberg 2012) with parameters: “--very-sensitive-local --no-discordant --no-mixed”. The resulting BAM files were parsed using Bedtools suite v2.25.0 utilities (Quinlan and Hall 2010). A genome file was created from BAM files using SAMtools view v1.3.1 (Danecek et al. 2021) and was divided into 50 kbp sliding windows by makewindows option with following parameters: “-w 50000 -s 50000”. The BAM files were sorted with SAMtools sort and converted to BED format using Bedtools bamtobed “-split”. We computed the coverage of aligned sequences in 50 kbp windows using Bedtools coverage and excluded reads which mapped to regions corresponding to repetitive sequences identified by RepeatModeler using the Bedtools subtract command. Finally, the median coverage depth for each 50 kbp window was computed with Bedtools genomecov. In both sexes, coverage depths for each scaffold were normalized by mean coverage across 50 kbp windows and compared between sexes, formulated as the Log2 of the male:female (M:F) coverage ratio. The resulting data were visualised together with the data from Pool-seq analysis using “SexGenomicsToolkit/sgtr” R package (https://github.com/SexGenomicsToolkit/sgtr).

## Results

### Genome size

We measured 2C values of nine blue butterfly species using flow cytometry. Resulting genome sizes with mean and standard deviation are shown in Table I along with the published karyotypes. The genome size of tested species ranges from 0.62 pg to 0.82 pg, with species with high chromosome number having larger genomes. However, the difference in genome size is no more than 32 %. Notably, Czech and Spanish populations of *P*. *icarus* differ in their genome sizes. According to these results we can conclude that increase in chromosome number in Lycaenidae did not result from polyploidization but rather from chromosomal fissions.

**Table I:** Male haploid DNA content (1C) of species under study determined by flow cytometry.

| Species | Specimen 1<br>(pg) | Specimen 2<br>(pg) | Specimen 3<br>(pg) | Mean 1C<br>(pg) | SD | 2n |
| --- | --- | --- | --- | --- | --- | --- |
| *P. icarus* (CZ) | 0.63 | 0.63 | 0.63 | 0.63 | 0.004 | 46<sup>a</sup> |
| *P. icarus* (ESP) | 0.62 | 0.62 | 0.62 | 0.62 | 0.001 | 46<sup>a</sup> |
| *P. escheri* (ESP) | 0.70 | 0.69 | 0.70 | 0.70 | 0.013 | 46<sup>b</sup> |
| *P. thersites* (ESP) | 0.67 | 0.66 | 0.67 | 0.66 | 0.012 | 48<sup>c</sup> |
| *L. bellargus* (CZ) | 0.67 | 0.65 | 0.66 | 0.66 | 0.008 | 90<sup>a</sup> |
| *L. hispana* (ESP) | 0.70 | 0.70 | 0.71 | 0.70 | 0.009 | 168<sup>d</sup> |
| *L. coridon* (CZ) | 0.71 | 0.69 | 0.71 | 0.70 | 0.015 | 179<sup>e*</sup> |
| *P. ripartii* (ESP) | 0.73 | 0.73 | 0.74 | 0.74 | 0.011 | 180<sup>f</sup> |
| *P. fulgens* (ESP) | 0.73 | 0.67 | 0.68 | 0.69 | 0.026 | 218<sup>f</sup> |
| *P. dorylas* (ESP) | 0.82 | n/a | n/a | 0.82 | n/a | 298<sup>g</sup> |

SD – Standard deviation; n/a – Missing values; \*Mean value of 2n = 174–184; <sup>a</sup>Nguyen et al. (2010); <sup>b</sup>de Lesse (1953) cited in Robinson (1971); <sup>c</sup>Lorković (1941) cited in Kandul et al. (2004); <sup>d</sup>Descimon and Mallet (2010); <sup>e</sup>de Lesse (1969) cited in Schmitt and Seitz (2001); <sup>f</sup>Munguira et al. (1995); <sup>g</sup>(Wiemers 2003).

### Cytogenetic analyses

We tried to determine the chromosome number of five species, namely *P*. *icarus*, *P*. *escheri*, *L*. *bellargus*, *L*. *coridon* and *P*. *dorylas*. The diploid chromosome number was counted from mitotic chromosomal preparations, which were stained by DAPI in DABCO. We confirmed the diploid chromosome number 2n = 46 in *P*. *icarus* males (Fig. S1d, f), while in females the number differed between individuals from 2n = 45 to 2n = 46 (Fig. 2a-b, S1c, e). We were not able to reliably count chromosomes in *P. escheri* due to the lack of chromosomal preparations, although two overlapping mitotic complements (Fig. 2c) seem to support the published chromosome number 2n = 46 (de Lesse 1953 cited in Robinson 1971). In case of specimens with greatly fragmented karyotype we were not able to infer their chromosomal number from our spread nuclei. We thus rely on chromosome numbers known from literature (Table I), although these belong to specimens from different populations.

**Figure 2.**
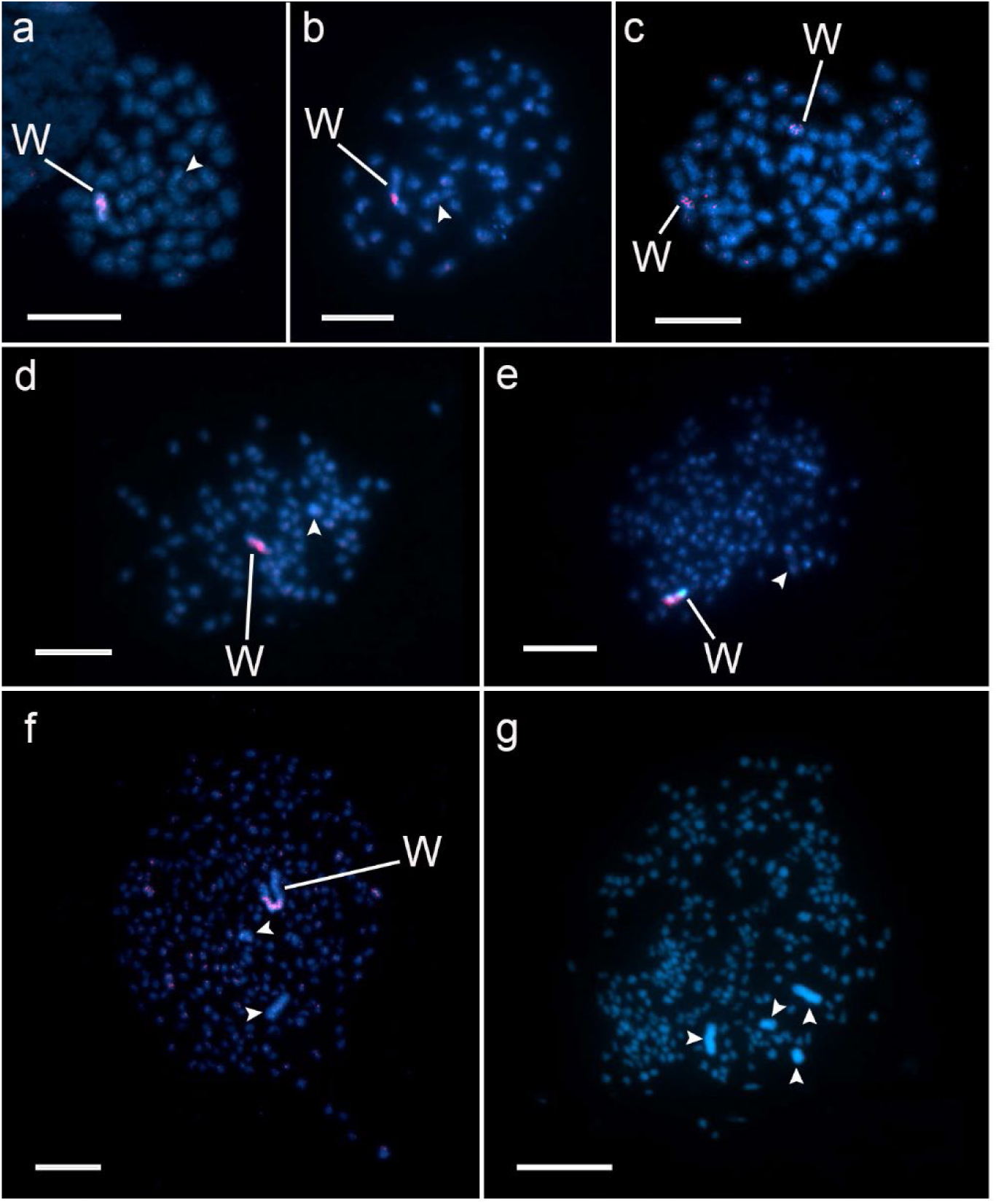
Genomic *in situ* hybridization on mitotic chromosome complements of blue butterflies under study. DAPI staining in blue and hybridization signals of female gDNA probes labelled with Cy3-dUTP in red. The arrowheads indicate putative chromosome Z identified according to its large sizes. (**a-b**) Mitotic nuclei of two *P. icarus* females which differ in their chromosome number. (**a**) Female with the karyotype 2n = 46 in which the female gDNA probe labelled a half of the W chromosome and (**b**) female with 2n = 45 in which GISH highlighted only the interstitial block of the W chromosome. (**c**) Two overlapping female mitotic nuclei of *P. escheri* with female genomic probe highlighting two small W chromosomes. We counted 2n = 46. (**d**) Female mitotic nucleus of *L. bellargus*. The female genomic probe clearly highlighted most of the chromosome W. Sex chromosomes stand out by their size. (**e**) Female mitotic nucleus of *L. coridon*. Sex chromosomes are again the largest chromosomes in the complement with GISH highlighting about half of the W chromosome. (**f**) Female mitotic nucleus of *P. dorylas*. The female genomic probe highlighted large interstitial region of the W chromosome, which is the largest chromosome in the karyotype. Two more clearly non-homologous chromosomes stand out by their size and likely correspond to two Z chromosomes. (**g**) *P. dorylas* male mitotic nucleus stained by DAPI support the presence of two pairs of Z chromosome, large and small. Scale bar 10 µm.

Staining of female pachytene complements by 2.5% lactic acetic orcein enabled detection of sex chromosome bivalents through differential chromomere patterns (Fig. 3; cf. Traut 1976). W chromosome heterochromatin should be deeply stained by lactic acetic orcein and thus easily distinguishable from Z chromosome and autosomes. That was the case in *P. icarus* (Fig. 3a–b) and *P. escheri* (Fig. 3c). In the other species, namely *L*. *bellargus*, *L*. *coridon* and *P*. *dorylas*, the difference was not so profound (Fig. 3d–f), though it was still noticeable. In *P. dorylas*, LAO highlighted only part of W sex chromosome (Fig. 3f). In *P. icarus* females with 2n = 46 or 2n = 45 LAO stained only a half or two small interstitial blocks, respectively (Fig. 3a–b). Pachytene nuclei of *P. escheri* comprise a small highly heterochromatic chromosome W which was deeply stained along its whole length. In most *P. escheri* nuclei we observed peculiar sex chromosomes pairing with larger Z chromosome wrapping around the smaller W chromosome (Fig. 3c).

**Figure 3.**
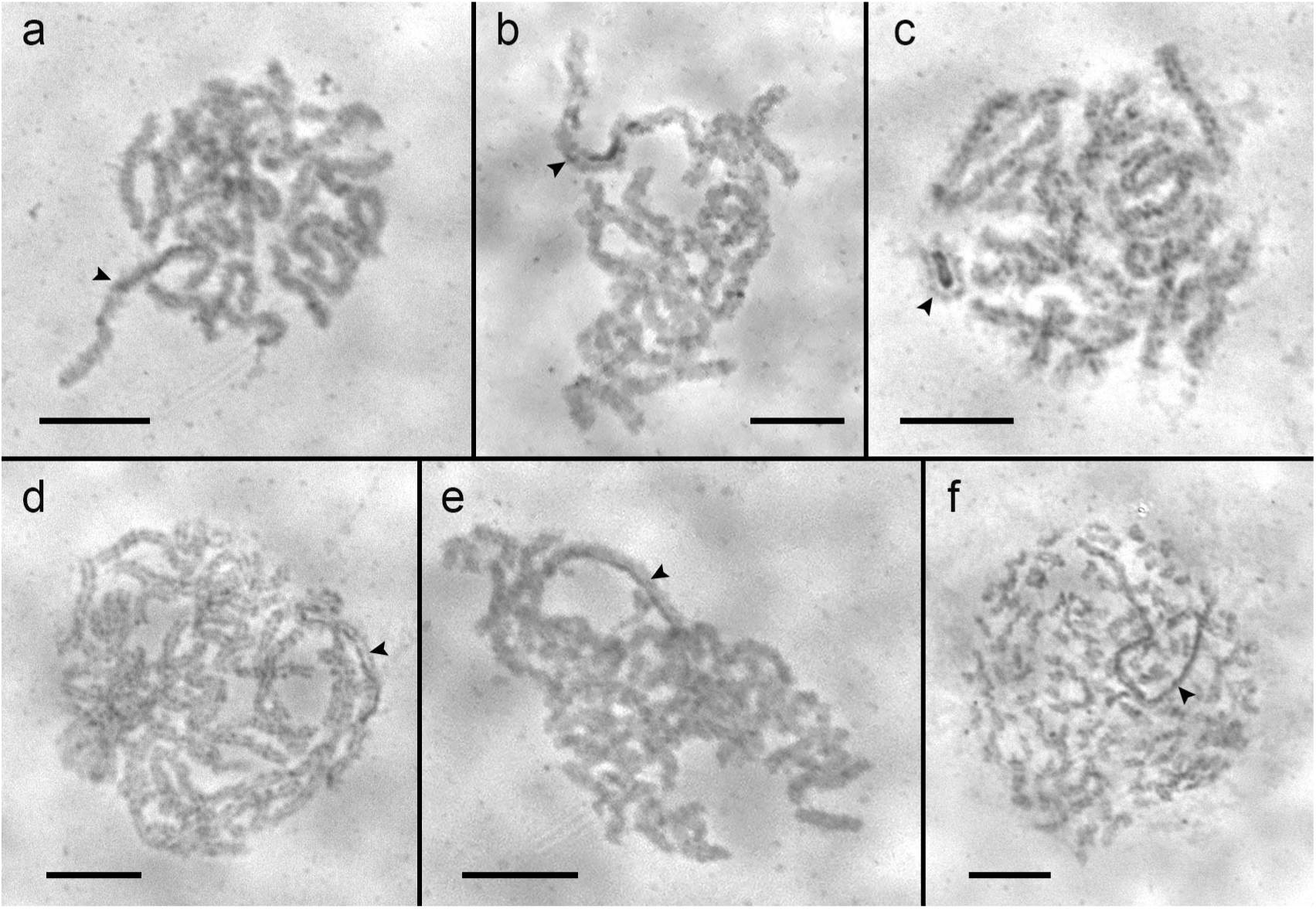
WZ pachytene bivalents detected by staining by 2.5% lactic acetic orcein in female nuclei. Arrowheads indicate sex chromosome bivalent distinguished by different chromomere pattern. (**a**) Female nucleus of *P*. *icarus* female with 2n = 46 where half part of W sex chromosome was highlighted by orcein. (**b**) Nucleus of *P*. *icarus* female with 2n = 45 and deeply stained interstitial blocks on the W chromosome. (**c**) *P*. *escheri* nucleus with the deeply stained W chromosome and characteristic pairing with the Z chromosome. (**d,e**) *L*. *bellargus* and *L. coridon* nuclei, respectively, with the W chromosome with weaker but still distinct pattern after LAO staining. (**f**) *P*. *dorylas* nucleus paired sex chromosomes are notably larger than autosomal bivalents. Lactic acetic orcein stained only a part of the W chromosome. Scale bar 10 µm.

For reliable detection of chromosome W, we used genomic *in situ* hybridization (GISH) on female spread chromosome preparations (Fig. 2 and 4). When compared to chromomere pattern after staining with LAO, GISH signals more less highlighted the same regions of W chromosomes. GISH confirmed that the sex chromosomes are the largest elements in all tested species except for *P*. *escheri*. Mitotic nuclei of females of *P*. *icarus*, *L. bellargus* and *L. coridon* carry two large sex chromosomes. Interestingly, in female mitotic nuclei of *P*. *dorylas* we observed three large chromosomes with the W chromosome being markedly larger than the other two (Fig. 2f), while in male mitotic nuclei we observed two large chromosome pairs (Fig. 2g). This implies that the largest W chromosome pairs with two Z chromosomes thus constituting WZ_1_Z_2_ multiple sex chromosome system (Fig. 2f).

**Figure 4.**
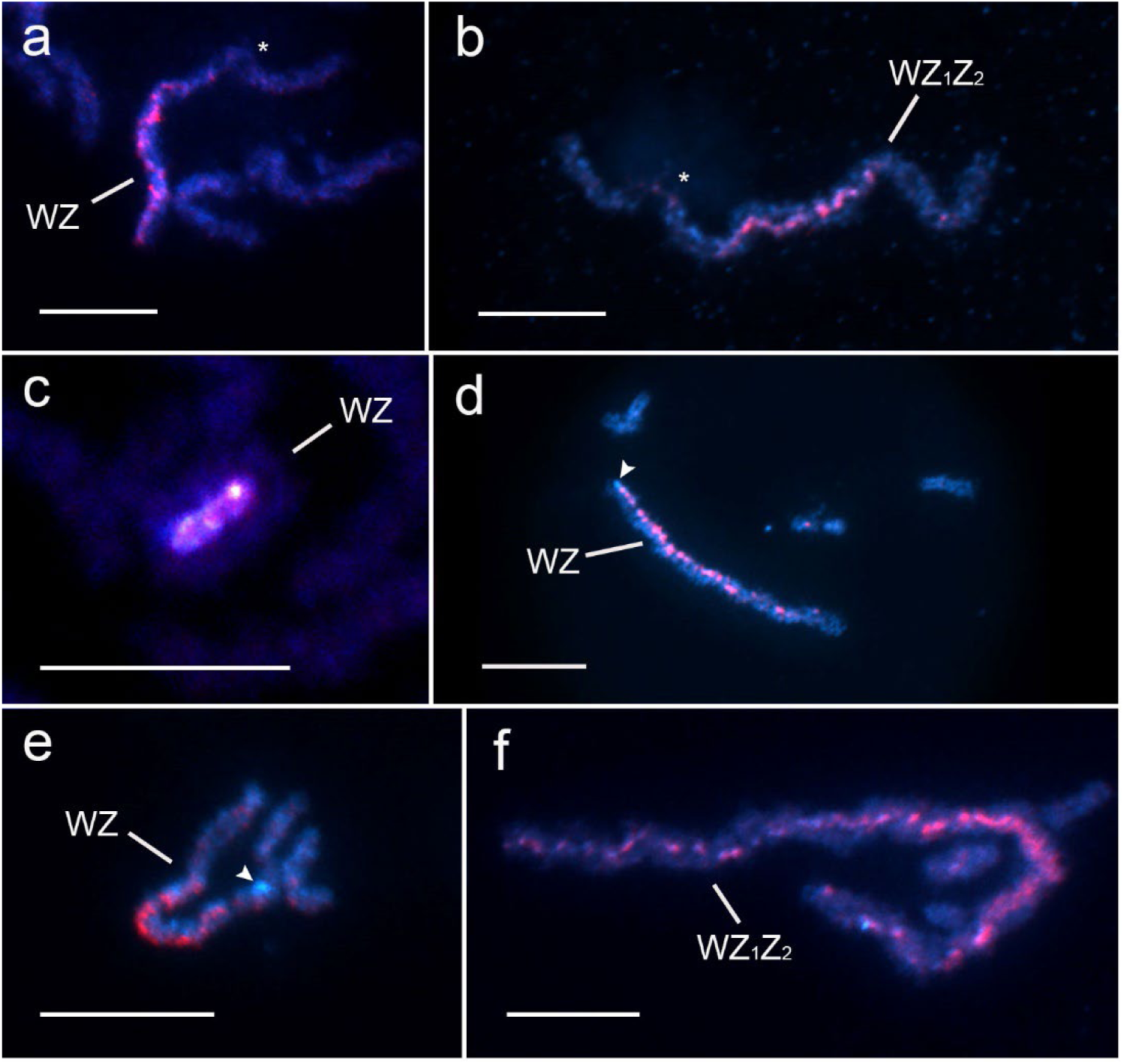
Genomic *in situ* hybridization on female pachytene chromosomes. Chromosomes were stained by DAPI (blue) and female gDNA probe was labelled with Cy3-dUTP or fluorescein-dUTP (red). Asterisks mark chromatin emanating into nucleolus. Arrowheads point to the DAPI positive heterochromatic blocks (**d**, **e**). (**a**) Sex chromosome bivalent of *P. icarus* with 2n = 46 with the probe labelling a half of the W chromosome. (**b**) Sex chromosomes in *P*. *icarus* with 2n = 45 and the probe hybridizing solely to the interstitial part of the W chromosome. In both karyotypic races we observed nucleolus associated with one of euchromatic regions of the W chromosome. (**c**) Sex chromosome bivalent in *P*. *escheri* with the probe highlighting the whole length of the W chromosome. Note characteristic pairing with the sex chromosome Z wrapping around the W. (**d**) Sex chromosome bivalent of species *L*. *coridon*. Note DAPI positive heterochromatic block at the end of this bivalent unlabelled by female genomic probe. Also, part of the opposite half bears only few interspersed signals. (**e**) Sex chromosome bivalent of *L*. *bellargus* with the hybridization pattern similar to *L. coridon* with weakly labelled DAPI postitive heterochromatin block, well differentiated strongly highlighted interstitial region and unlabelled region. (**f**) Paired sex chromosomes of *P*. *dorylas* with the probe strongly highlighting the interstitial part of W chromosome and weaker interspersed signals covering the rest of the W chromosome. Scale bar 10 µm.

As mentioned above, in *P*. *icarus* the W sex chromosomes varied between females with different chromosome numbers. In all tested specimens with 2n = 45 GISH labelled solely interstitial part of W chromosome in pachytene nuclei (Fig. 4b, 5b, S1b). The same pattern was observed also in mitotic nuclei (Fig. 2b, S1e) and after staining by LAO (Fig. 3b). In females with 2n = 46 GISH labelled half of the W chromosome (Fig. 2a, 4a, 5a, S1a, c) which implies the presence of karyotypic races which differ in their sex chromosome constitution. The results of GISH also revealed a considerable difference between W chromosomes of *P. icarus* (2n = 46) and *P. escheri* (Fig. 4a, 4c), although their chromosome number is the same (Tab. I). Unlike in *P. icarus*, the whole *P. escheri* W chromosome was highlighted by the female genomic probe. Moreover, we confirmed the peculiar pairing between the W and Z sex chromosomes (Fig. 4c, S1g). In both *L. coridon* and *L. bellargus* we observed DAPI positive block in one end of their sex chromosome bivalents which has not been labelled by female genomic probe. Despite LAO stained more less the whole length of their W chromosomes, the female genomic probe labelled only about two thirds of their W chromosomes in both species (Fig. 4d–e). In *P. dorylas*, the female genomic probe strongly hybridized mainly to interstitial part of W chromosome, similar to LAO staining pattern, with more interspersed signals along the rest of the W (Fig. 4f, 5f, S1j).

**Figure 5.**
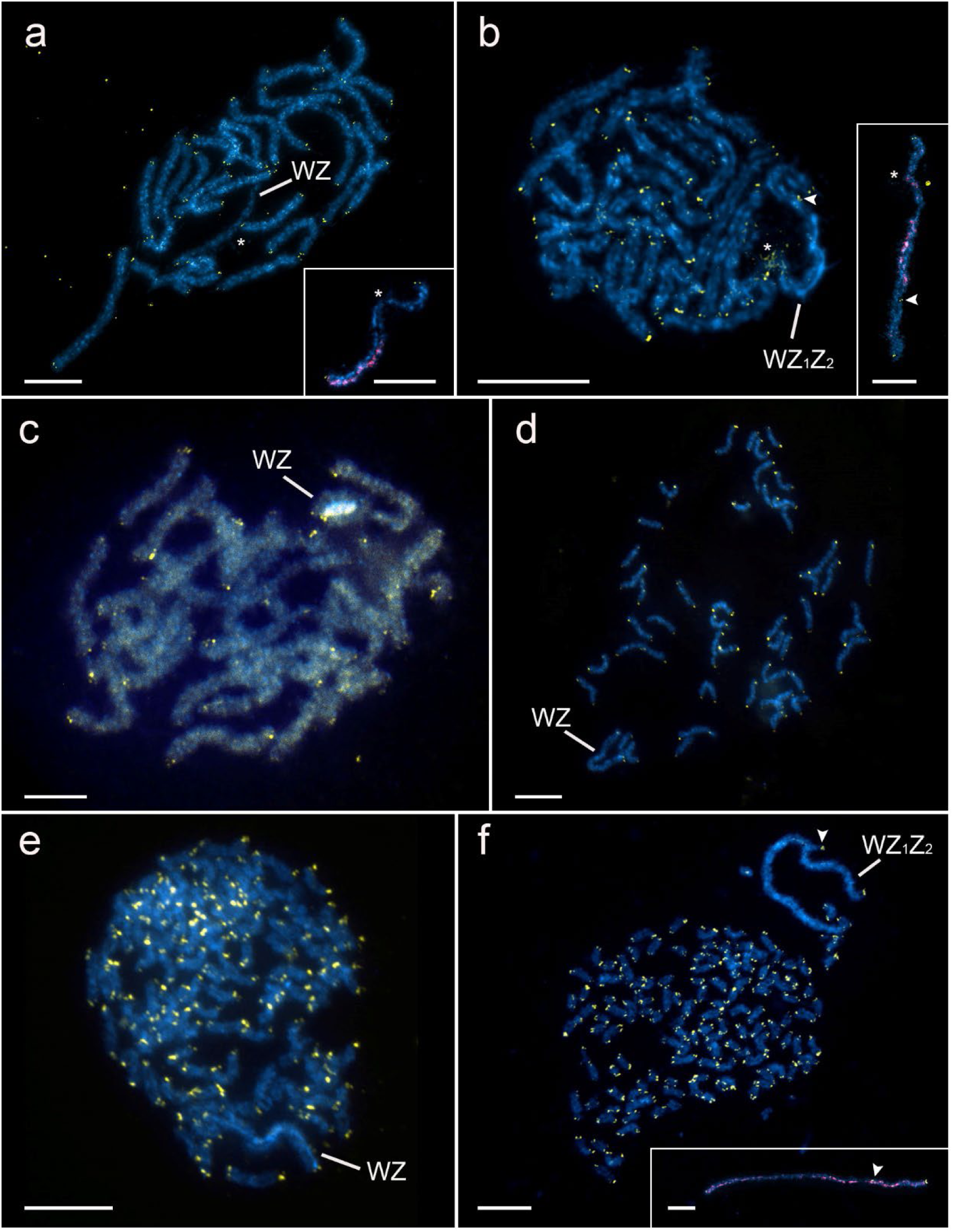
Fluorescence *in situ* hybridization with the (TTAGG)_n_ telomeric probe in females pachytene nuclei. Chromosomes were stained by DAPI (blue), the (TTAGG)_n_ probe was labelled with Cy3-dUTP or fluorescein-dUTP (yellow). Asterisks mark chromatin emanating into nucleolus. Arrowheads mark the interstitial telomeric signals. The telomeric probe was combined with GISH with female gDNA labelled by Cy3-dUTP or fluorescein dUTP (red) in insets in (**a**), (**b**) and (**f**). (**a**) *P*. *icarus* with 2n = 46, WZ, (**b**) *P*. *icarus* with 2n = 45, WZ_1_Z_2_, (**c**) *P. escheri*, (**d**) *L. bellargus*, (**e**) *L. coridon* and (**f**) *P. dorylas*. In all cases we observed signals of telomeric sequences at the ends of all chromosomes. Moreover, we also detected interstitial telomeric signals located in sex chromosome multivalents in one of two karyotypic races of *P*. *icarus* (**b**) and in *P*. *dorylas* (**f**). Scale bar 10 μm.

After GISH, chromosomal preparations were re-probed with (TTAGG)_n_ telomeric probe (telo-FISH) to test for the presence of telomeric sequences in species with high chromosome numbers and detection of sex chromosome multivalents.

In all tested species we confirmed presence of telomeric sequences in both ends of all chromosomes, which corroborates their *de novo* synthesis after fragmentation (Fig. 5). Moreover, we detected interstitial telomeric sequences (ITS) located in sex chromosome multivalents in two species, namely the *P. icarus* karyotypic race with 2n = 45 (Fig. 5b) and *P. dorylas* (Fig. 5f). In *P. icarus* the ITS was detected at the boundary of euchromatic part, which did not carry nucleolus organizer region (NOR), and interstitial heterochromatic part of W chromosome highlighted with female genomic probe (Fig. 5b). These findings together with difference in chromosome number between males and females (see above) imply multiple sex chromosome system in this karyotypic race of *P. icarus*. The presence of ITS located in sex chromosome multivalent in *P. dorylas* (Fig. 5f) agrees with observation of three non-homologous large chromosomes in female mitotic nuclei, which therefore correspond to chromosomes W, Z_1_ and Z_2_ (Fig. 2f). In the other species under study, namely *P. escheri* (Fig. 5c), *L. bellargus* (Fig. 5d) and *L. coridon* (Fig. 5e), we observe no ITS located in their paired sex chromosomes.

As it seems, sex chromosomes are associated with nucleolar organizing region (NOR) in *P. icarus* (Fig. 4a–b, 5a–b), we performed fluorescence *in situ* hybridization with the 18S rDNA probe (Fig. S1). FISH with the 18S rDNA probe confirmed sex-linked rDNA clusters in both *P. icarus* karyotypic races. In pachytene bivalents, the hybridization signal was localized at the border of euchromatic and heterochromatic W chromosome regions (Fig. S1a–b). However, the rDNA FISH on female and male mitotic complements revealed that the W sex chromosomes do not bear NOR. The rDNA cluster was detected on the two largest elements of mitotic complement corresponding to Z chromosomes in both races (Fig. S1c–f). Thus, we can infer that clusters of 18S rDNA genes are Z-linked in *P. icarus*. However, in *P. escheri* the rDNA probe hybridized to a single autosomal bivalent (Fig. S1g). Two hybridization signals of different intensities were localized on autosomal bivalents of *L. bellargus* (Fig. S1h). The probe hybridized only to three autosomes in mitotic complements, so we hypothesize that one of observed rDNA locus is heterozygous (not shown). In *L. coridon,* clusters of rDNA genes were located on two autosomal bivalents (Fig. S1i) which was confirmed by detection of four hybridization signals in mitotic complements (not shown). On the contrary, only one autosomal bivalent carried the major rDNA cluster in pachytene complements of *P. dorylas* (Fig. S1j), which is in accordance with two hybridization signals observed in mitotic nuclei.

### Pool-seq and coverage data analysis

We combined two different approaches to identify sex-specific regions, namely, pool-seq and coverage analysis as these are suitable for identification of sex chromosomes with both low and high level of differentiation (Palmer et al. 2019). All autosomes but chromosome 20 showed low Fst and no differences in coverage betweeen sexes or sex-specific SNPs (Fig. 6a, Fig. S2). While the chromosome 20 also showed no differences in coverage between sexes, it had windows with increased F_st_ and female-specific SNPs, which makes it candidate for the Z_2_ chromosome of recent fusion origin (Fig. 6b, Fig. S2). Our data confirmed that Z chromosome consist of two synteny blocs. Pool-seq analysis identified a region with increased F_st_ and female-specific SNPs, which has also low level of differentiation as suggested by no difference in coverage between sexes. This region thus represents a recent addition. The rest of the Z chromosome has clear differences in coverage without sexes. Since there is little similarity in this region, it has also low F_st_ and ammount of sex-specific SNPs. This region thus corresponds to the ancestral Lepidopteran Z chromosome (Fig. 6c, Fig. S2).

**Figure 6:**
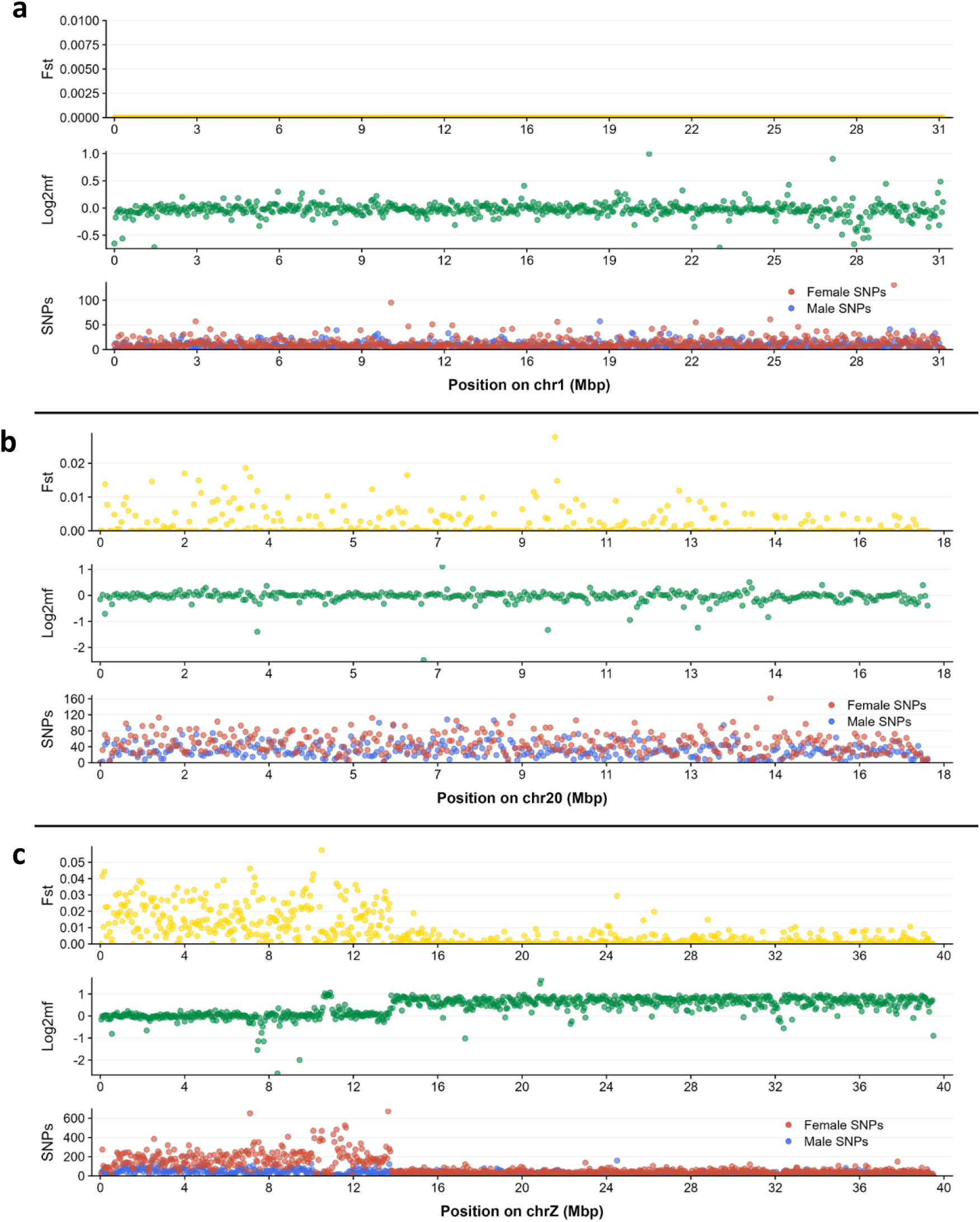
Identification of sex-linked regions in *P. icarus* race with WZ_1_Z_2_ sex chromosome constitution. Plot shows results of coverage (green) and Pool-seq analyses (male- and female-specific SNPs and their relative divergence - F_st_ are in blue, red and yellow, respectively). (**a**) Resulting values for the chromosome 1 which represents majority of autosomes. (**b**) Chromosome 20 with raised number of male- and female-specific SNPs but no difference in coverage between sexes probably corresponds to the Z_2_ chromosome. (**c**) Neo-Z chromosome comprising an ancestral region with clear difference in coverage between sexes and novel addition with no difference in coverage but higher number of female-specific SNPs and increased F_st_.

## Discussion

Based on two- and fourfold differences in chromosome numbers observed between closely related lycaenid species such as *P. icarus* (n = 22–23), *L. bellargus* (n = 45), and *L. coridon* (n = 87–97), polyploidy was hypothesized as a possible mode of karyotype evolution in these butterflies. Polyploidy in lycaenids has been previously refuted by indirect evidence such as comparison of areas occupied by native metaphase I plates or negative correlarion between number of chromosomes and their size (Kandul et al. 2007, Pazhenkova and Lukhtanov 2023). More recently, length of genome assemblies as well as gene syntenies favored chromosome fragmentation over whole genome duplications in *L. bellargus and L. coridon* (Lohse et al. 2022, Vila et al. 2023, Pazhenkova and Lukhtanov 2023, Wright et al. 2024). Our results provide first genome size values determined by flow cytometry for representatives of all three polyommatine lineages with high chromosome numbers. The results do not provide any support for polyploidy in the polyommatine butterflies, although the species with high chromosome numbers seem to have larger genomes than their outgroups with ancestral karyotype (Fig. 1, Table I). This is in contrast with hypothesis of holocentric drive which should promote either chromosomal fusions and/or multiplication of transposable elements (TEs) when larger chromosomes preferentially segregate to oocytes in female asymetric meiosis or chromosomal fissions and/or depletion of TEs when smaller chromosomes are favoured, thus producing negative correlation between chromosome number and genome size (Bureš and Zedek 2014). Indeed, a recent study in the *Leptidea* wood whites suggests that meiotic drive does not promote karyotype evolution but rather opposes chromosome fusions and stabilizes lepidopteran karyotypes (Boman et al. 2024).

While the chromosome-level assemblies of *L. bellargus*, *L. coridon*, and *P. icarus* were 0.54 pg, 0.55 pg and 0.52 pg, respectively (Lohse et al. 2022, Lohse and Vila 2023, Vila et al. 2023), flow cytometry estimated the genome sizes to be much higher with 0.66 pg, 0.70 pg and 0.63 pg, respectively. Flow cytometry is considered a more reliable approach compared to sequence-based methods (Pflug et al. 2020). So it seems even genome assemblies built using long reads underestimate genome sizes (cf. Mora et al. 2024). Providing that differences in genome sizes are determined by repetitive elements (Pazhenkova and Lukhtanov 2023), increase in genome sizes in species with fragmented karyotypes supports an instrumental role of mobile elements in karyotypic diversification of *Lysandra* and *Polyommatus* spp. However, even in our small sample, we also observed considerable differences in genome sizes between and within populations in *P. icarus* and *P. fulgens* (Table I). Indeed, study of repeat landscapes in *Erebia* butterflies showed that repeat composition can vary greatly within species and the variation scales with population divergence (Cornet et al. 2023). Further study is needed to confirm whether increase in genome size can be ascribed to activity of horizontally transferred mobile elements (Kandul et al. 2004).

It was hypothesized, that interstitial telomeric sequences (ITS) serve as fission breakpoints facilitating the fragmentation of chromosomes as they can induce and heal chromosome breaks (Pazhenkova and Lukhtanov 2023). In polyommatine butterflies, telomeres have been so far examined by means of molecular cytogenetics only in several representatives of the subgenus *Agrodiaetus* with chromosome numbers ranging from n=10 to n=ca. 108 (Vershinina et al. 2015). However, telomeric signals were weak and difficult to distinguish from background noise especially in species with high chromosome numbers, supposedly due to small size of mitotic chromosomes at the limit of resolution of light microscopy (Vershinina et al. 2015). Pachytene bivalents provided much better resolution revealing clear telomeric signals at the ends of chromosomes even in *Lysandra* spp. and *P.* (subgenus *Plebicula*) *dorylas* with high chromosome numbers, which seem to display even stronger signals then species with modal chromosome numbers (Fig. 5). In the chromosome level assemblies of *L. bellargus* and *L. coridon*, telomeric repeats were not found in about 10 % of chromosomes (Pazhenkova and Lukhtanov 2023). Our results support a hypothesis that these pseudochromosomes are truncated as we found clear telomeric signals at the ends of nearly all chromosomes (Fig. 5d, e). Interestingly, compared to their relatives with modal karyotypes telomeric signals seem to be stronger in *L. coridon and P. dorylas* with high chromosome numbers (Fig. 5e, f). This suggests increased activity of telomerase, which could heal chromosome ends by synthesizing telomeres de novo. Indeed, in *B. mori*, it was shown that ends of chromosome fragments induced by X-ray irradiation were healed by de novo addition of (TTAGG)_n_ repeats, which are important for their stable transmision (Fujiwara et al. 2000). We failed to reliably detect any autosomal interstitial telomeric sequences reported in *L. bellargus* and *L. coridon* (Pazhenkova and Lukhtanov 2023), which is not surprising as the length of these ITS seems to be <1.5 Kb (Pazhenkova and Lukhtanov 2023), which is below detection threshold of conventional FISH protocols (Speel et al. 2000, Carabajal Paladino et al. 2014).

However, we detected interstitial telomeric signals on paired sex chromosomes of *P. icarus* females with 2n = 45 (Fig. 5b) and *P. dorylas* females (Fig. 5f). Differences in chromosome numbers between sexes in *P. icarus* as well as presence of additional euchromatic region on the W chromosome (Fig. 4a, b) suggest, that the interstitial signals do not correspond to ITS but rather indicate sex chromosome trivalent WZ_1_Z_2_ originating from sex chromosome-autosome fusions (cf. Yoshido et al. 2005). Such neo-sex chromosome systems are common in Lepidoptera (e.g. Carabajal Paladino 2019, Mora et al. 2024, Wright et al. 2024). Four independent Z-autosome fusions have been reported from chromosome level assemblies of several lycaenids including *L. coridon* and *P. icarus*. The results suggest that the reported *P. icarus* fusion was formed between the Z chromosome and an autosome bearing arrays of rDNA, which is Z-linked in *P. icarus* females with both 2n = 45 and 2n = 46 (Fig. S1a–f). We assume a corresponding fusion occured between the W chromosome and the autosome resulting in a neo-W chromosome with one euchromatic region (Fig. 2a, 4a) and possibly involving loss of the rDNA cluster. This neo-sex chromosome system seems to be wide spread as it is shared between Scottish (Lohse and Vila 2023) and Czech *P. icarus* populations and thus could be up to 2 My old (cf. Dinca et al. 2011). However, another fusion occured only between the neo-W chromosome and another autosome in the Czech population giving rise to the observed multiple sex chromosome system with yet another euchromatic region on the W chromosome and chromosome Z_2_ (Fig. 4a–b, 5a–b). Thus, we observed two distinct karyotype races in *P. icarus.* Genomic analyses identified the chromosome 20 as a candidate for Z_2_ chromosome (Fig. 6b, Fig. S2). Repeated sex chromosome-autosome fusions resulting in two races make *P. icarus* ideal system for future studies of drivers of sex chromosome turnovers (cf. Mora et al. 2024).

Similarly, sex-linked interstitial telomeric signals (Fig. 5f) and different constitutions of large chromosomes in male and female mitotic complements (Fig. 2f, g) suggest presence of the multiple sex chromosome systems WZ_1_Z_2_ in *P. dorylas*. The female mitotic complements comprised three distinct large chromosomes, the largest being W chromosome (Fig. 2f), while male karyotype contained two large pairs (Fig. 2f–g). The hybridization pattern of the female genomic probe in pachytene nuclei suggested that the *P. dorylas* W chromosome comprises a highly differentiated interstitial region and two less differentiated regions (Fig. 4f), which could again indicate presence of two independent W chromosome-autosome fusions. Interestingly, the interstitial telomeric signal was located in a region pairing with the highly differentiated part of the W sex chromosome (Fig. 5f), which would suggest fission rather than fusion origin of the Z_2_ chromosome in *P. dorylas*. Futher study is needed to discern between these two scenarios. Two large chromosomal pairs were observed also in other species of subgenus *Plebicula*, namely *Polyommatus golgus* (2n = 268), *P. nivescence* (2n = 382), and *P. atlantica* (2n = 446) (de Lesse 1970, Lukhtanov 2015), which suggests that the multiple sex chromosome system occured in a common ancestor of this lineage. Accordingly, scattered GISH hybridization signals indicate higher level of differentiation of these W chromosome regions (Fig. 4f).

Sex chromosomes are clearly the largest elements in karyotypes of *L. bellargus*, *L. coridon* and *P. dorylas* (Fig. 2d–g) and must have resisted extensive chromosome fragmentation. Chromosome Z avoids fissions also in other lepidopteran lineages with highly reorganized genomes (Wright et al. 2024), although it is generally involved in highest number of fusions (Carabajal Paladino et al. 2019, Wright et al. 2024) and also experiences much more intrachromosomal rearrangments compared to autosomes (Van’t Hof et al. 2013). Similarly, an aphid X chromosome seems to be highly conserved, in this case, however, even recalcitrant to rearrangements with autosomes (Mathers et al. 2021). Also gene content of a mammalian X chromosome was early recognized as evolutionarily highly conserved by Susumu Ohno (1967). Ohno (1967) proposed that it is constrained by dosage compensation. More recently, it was suggested that the strong selective constraint is actually imposed by the 3D genomic architecture needed for X chromosome inactivation (Brashear et al. 2021). Moreover, intact aphid X chromosomes may be a must for proper elimination of the X chromosome during male sex determination (Mathers et al. 2021).

However, lepidopteran sex chromosomes can be broken. Species-specific sex chromosome fissions occurred in *Leptidea* wood white butterflies, yet the resulting elements can be still paired into complex multivalents, which segregate properly due to achiasmatic and inverted female meiosis (Lukhtanov et al. 2018, Yoshido et al. 2020). The non-homologous W and Z chromosomes can pair despite considerable size differences through synaptic adjustment (Marec 1996, Van’t Hof et al. 2008). Moreover, missegregation of W chromosome fragments without sex determining genes should bear no fitness cost as shown in the *Samia* wild silkmoths (Yoshido et al. 2016). So it is particularly intriguing that the W chromosome is structurally the most conserved chromosome in polyommatine karyotypes. This suggest the W chromosome to be protected from fissions rather than selectively constrained. Indeed, the *B. mori* W chromosome is epigenetically repressed in both somatic tissues and germline except for postpachytene (Traut et al. 2019) and expresses piRNAs supressing transposable elements (Kawaoka et al. 2011, Hara et al. 2012) and possibly acting in cis (cf. Gebert et al. 2021). We hypothesize that highly repetitive but epigeneticly silenced W chromosome is protected from fisions caused by mobile elements and its pairing with Z chromosomes or their fragments ensures their proper segregation and allows for fusions of their sticky ends, thus contributing to structural integrity to the Z chromosome. In general, it seems that evolutionary stability of sex chromosomes can be a convergent feature resulting from various idiosyncracies of sex chromosomes.

## Acknowledgement

This research was funded by the grant 20-20650Y of the Czech Science Foundation. G.T. was supported by the grant PID2020-117739GA-I00 MCIN/AEI/10.13039/ 501100011033 and by the grant 2021-SGR-01334 from the Departament de Recerca i Universitats (Generalitat de Catalunya). Computational resources were provided by the e-INFRA CZ project (ID:90254), supported by the Ministry of Education, Youth and Sports of the Czech Republic, and by the ELIXIR-CZ project (ID:90255), part of the international ELIXIR infrastructure.

## References

Andrews, S. (2010). FastQC: A quality control tool for high throughput sequence data. https://www.bioinformatics.babraham.ac.uk/projects/fastqc/

Augustijnen, H., Patsiou, T., Schmitt, T., & Lucek, K. (2024). Living on the Edge—Genomic and Ecological Delineation of Cryptic Lineages in the High-Elevation Specialist *Erebia Nivalis*. Insect Conservation and Diversity, 17(3), 526–542. 10.1111/icad.12721

Bolger, A. M., Lohse, M., & Usadel, B. (2014). Trimmomatic: a flexible trimmer for Illumina sequence data. Bioinformatics, 30(15), 2114–2120. 10.1093/bioinformatics/btu170

Boman, J., Wiklund, C., Vila, R., & Backström, N. (2024). Meiotic drive against chromosome fusions in butterfly hybrids. Chromosome Research, 32(2), 7. 10.1007/s10577-024-09752-0

Brashear, W. A., Bredemeyer, K. R., & Murphy, W. J. (2021). Genomic architecture constrained placental mammal X Chromosome evolution. Genome Research, 31(8), 1353–1365. 10.1101/gr.275274.121

Buntrock, L., Marec, F., Krueger, S., & Traut, W. (2012). Organ Growth without Cell Division: Somatic Polyploidy in a Moth, *Ephestia Kuehniella*. Genome, 55(11), 755–763. 10.1139/g2012-060

Bureš, P., & Zedek, F. (2014). Holokinetic Drive: Centromere Drive in Chromosomes without Centromeres. Evolution, 68(8), 2412–2420. 10.1111/evo.12437

Carabajal Paladino, L. Z., Nguyen, P., Šíchová, J., & Marec, F. (2014). Mapping of single-copy genes by TSA-FISH in the codling moth. BMC Genomics, 15(S15). 10.1186/1471-2156-15-S2-S15

Carabajal Paladino, L. Z., Provazníková, I., Berger, M., Bass, C., Aratchige, N. S., López, S. N., Marec, F., & Nguyen, P. (2019). Sex Chromosome Turnover in Moths of the Diverse Superfamily Gelechioidea. Genome Biology and Evolution, 11(4), 1307–1319. 10.1093/gbe/evz075

Carey, S. B., Lovell, J. T., Jenkins, J., Leebens-Mack, J., Schmutz, J., Wilson, M. A., & Harkess, A. (2022). Representing Sex Chromosomes in Genome Assemblies. Cell Genomics, 2(5), 100132. 10.1016/j.xgen.2022.100132

Cornet, C., Mora, P., Augustijnen, H., Nguyen, P., Escudero, M., & Lucek, K. (2023). Holocentric repeat landscapes: From micro-evolutionary patterns to macro-evolutionary associations with karyotype evolution. *Molecular Ecology*, Online ahead of print. 10.1111/mec.17100

Danecek, P., Bonfield, J. K., Liddle, J., Marshall, J., Ohan, V., Pollard, M. O., Whitwham, A., Keane, T., McCarthy, S. A., Davies, R. M., & Li, H. (2021). Twelve years of SAMtools and BCFtools. GigaScience, 10, 1–4. 10.1093/gigascience/giab008

Descimon, H., & Mallet, J. (2010) Bad Species. In J. Settele, T. G. Shreeve, M. Konvička, & V. H. Dyck (Ed.), Ecology of Butterflies in Europe (pp. 219-249). Cambridge, Cambridge University Press.

Drinnenberg, I. A., deYoung, D., Henikoff, S., & Malik, H. S. (2014). Recurrent Loss of *CenH3* Is Associated with Independent Transitions to Holocentricity in Insects. ELife, 3(e03676). 10.7554/eLife.03676

Ellegren, H. (2011). Sex-chromosome evolution: recent progress and the influence of male and female heterogamety. Nature Reviews Genetics, 12(3), 157–166. 10.1038/nrg2948

Ennis, T. J. (1976). Sex Chromatin and Chromosome Numbers in Lepidoptera. Canadian Journal of Genetics and Cytology, 18(1), 119–130. 10.1139/g76-017

Escudero, M., Márquez-Corro, J. I., & Hipp, A. L. (2016). The Phylogenetic Origins and Evolutionary History of Holocentric Chromosomes. Systematic Botany, 41(3), 580–585. 10.1600/036364416X692442

Faria, R., & Navarro, A. (2010). Chromosomal Speciation Revisited: Rearranging Theory with Pieces of Evidence. Trends in Ecology and Evolution, 25(11), 660–669. 10.1016/j.tree.2010.07.008

Fuková, I., Nguyen, P., & Marec, F. (2005). Codling Moth Cytogenetics: Karyotype, Chromosomal Location of RDNA, and Molecular Differentiation of Sex Chromosomes. Genome, 48(6), 1083–1092. 10.1139/g05-063

Fujiwara, H., Nakazato, Y., Okazaki, S., & Ninaki, O. (2000). Stability and Telomere Structure of Chromosomal Fragments in Two Different Mosaic Strains of the Silkworm, *Bombyx mori*. Zoological Science, 17(6), 743–750. 10.2108/zsj.17.743

Gebert, D., Neubert, L. K., Lloyd, C., Gui, J., Lehmann, R., & Teixeira, F. K. (2021). Large *Drosophila* germline piRNA clusters are evolutionarily labile and dispensable for transposon regulation. Molecular Cell, 81(19), 3965–3978.e5. 10.1016/j.molcel.2021.07.011

Hara, K., Fujii, T., Suzuki, Y., Sugano, S., Shimada, T., Katsuma, S., & Kawaoka, S. (2012). Altered expression of testis-specific genes, piRNAs, and transposons in the silkworm ovary masculinized by a W chromosome mutation. BMC Genomics, 13(19). 10.1186/1471-2164-13-119

Kandul, N. P., Lukhtanov, V. A., Dantchenko, A. V., Coleman, J. W. S., Sekercioglu, C. H., Haig, D., & Pierce, N. E. (2004). Phylogeny of Agrodiaetus Hübner 1822 (Lepidoptera: Lycaenidae) Inferred from MtDNA Sequences of COI and COII and Nuclear Sequences of EF1-α: Karyotype Diversification and Species Radiation. Systematic Biology, 53(2), 278–98. 10.1080/10635150490423692

Kandul, N. P., Lukhtanov, V. A., & Pierce, N. E. (2007). Karyotypic Diversity and Speciation in *Agrodiaetus* Butterflies. Evolution, 61(3), 546–559. 10.1111/j.1558-5646.2007.00046.x

Kawaoka, S., Kadota, K., Arai, Y., Suzuki, Y., Fujii, T., Abe, H., Yasukochi, Y., Mita, K., Sugano, S., Shimizu, K., Tomari, Y., Shimada, T., & Katsuma, S. (2011). The silkworm W chromosome is a source of female-enriched piRNAs. RNA, 17(12), 2144–2151. 10.1261/rna.027565.111

Langmead, B., & Salzberg, S. L. (2012). Fast gapped-read alignment with Bowtie 2. Nature Methods, 4(9), 357–359. 10.1038/nmeth.1923

de Lesse, H. (1970). Les Nombres de Chromosomes Dans Le Groupe de *Lysandra Argester* et Leur Incidence Sur Sa Taxonomie [Lep. Lycaenidae]. Bulletin de la Société entomologique de France, 75, 64–68. 10.3406/bsef.1970.21116

Li, H., & Durbin, R. (2009). Fast and accurate short read alignment with Burrows–Wheeler transform. Bioinformatics, 25(14), 1754–1760. 10.1093/bioinformatics/btp324

Lohse, K., Hayward, A., & Vila, R. (2022). The genome sequence of the Adonis blue, *Lysandra bellargus* (Rottemburg, 1775). Wellcome Open Research, *12*(7), 255. 10.12688/wellcomeopenres.18330.1

Lohse, K., & Vila, R. (2023). The genome sequence of the Common Blue, *Polyommatus icarus* (Rottemburg, 1775). Wellcome Open Research, 8(72). 10.12688/wellcomeopenres.18772.1

Lucek, K., Augustijnen, H., & Escudero, M. (2022). A Holocentric Twist to Chromosomal Speciation? Trends in Ecology & Evolution, 37(8), 655–662. 10.1016/j.tree.2022.04.002

Lukhtanov, V. A. (2015). The Blue Butterfly Polyommatus (Plebicula) Atlanticus (Lepidoptera, Lycaenidae) Holds the Record of the Highest Number of Chromosomes in the Non-Polyploid Eukaryotic Organisms. Comparative Cytogenetics, 9(4), 683–690. 10.3897/CompCytogen.v9i4.5760

Lukhtanov, V. A., Dantchenko, A. V., & Kandul, N. P. (1997). Die Kaiyotypen von Polyommatus (Agrodiaetus) Damone Damone Und P. (A.) Da Modes Rossicus Nebst Einigen Problemen Bei *Agrodiaetus* (Lepidoptera: Lycaenidae). Nachrichten Des Entomologischen Vereins Apollo, 16, 43–48. 10.14411/eje.2017.023

Lukhtanov, V. A., Dincă, V., Friberg, M., Šíchová, J., Olofsson, M., Vila, R., Marec, F., & Wiklund, C. (2018). Versatility of multivalent orientation, inverted meiosis, and rescued fitness in holocentric chromosomal hybrids. Proceedings of the National Academy of Sciences, 115(41), E9610–E9619. 10.1073/pnas.1802610115

Marec, F. (1990). Genetic control of pest Lepidoptera: Induction of sex-linked recessive lethal mutations in *Ephestia kuehniella* (Pyralidae). Acta Entomologica Bohemoslovaca, 87, 445–458.

Marec, F. (1996). Synaptonemal complexes in insects, International Journal of Insect Morphology and Embryology. International Journal of Insect Morphology and Embryology, 25(3), 205–233. 10.1016/0020-7322(96)00009-8

Martin, M. (2011). CUTADAPT removes adapter sequences from high-throughput sequencing reads. EMBnet Journal, 17(1). 10.14806/ej.17.1.200

Mathers, T. C., Wouters, R. H. M., Mugford, S. T., Swarbreck, D., van Oosterhout, C., & Hogenhout, S. A. (2021). Chromosome-Scale Genome Assemblies of Aphids Reveal Extensively Rearranged Autosomes and Long-Term Conservation of the X Chromosome. Molecular Biology and Evolution, 38(3), 856–875. 10.1093/molbev/msaa246

Mediouni, J., Fuková, I., Frydrychová, R., Dhouibi, M. H., & Marec, F. (2004). Karyotype, Sex Chromatin and Sex Chromosome Differentiation in the Carob Moth, *Ectomyelois Ceratoniae* (Lepidoptera: Pyralidae). Caryologia, 57, 184–194. 10.1080/00087114.2004.10589391

Mora, P., Hospodářská, M., Chung Voleníková, A., Koutecký, P., Štundlová, J., Dalíková, M., Walters, J. R., & Nguyen, P. (2024). Sex-Biased Gene Content Is Associated with Sex Chromosome Turnover in Danaini Butterflies. Molecular Ecology, 5(e17256). 10.1111/mec.17256

Munguira, M., Martin, J., & Pérez-Valiente, M. (1995). Karyology and Distribution as Tools in the Taxonomy of Iberian *Agrodiaetus* Butterflies (Lepidoptera : Lycaenidae). Nota Lepidopterologica, 17(3/4), 125–140.

Nguyen, P., Sahara, K., Yoshido, A., & Marec, F. (2010). Evolutionary Dynamics of RDNA Clusters on Chromosomes of Moths and Butterflies (Lepidoptera). Genetica, 138(3), 343–354. 10.1007/s10709-009-9424-5

Nguyen, P., Sýkorová, M., Šíchová, J., Kůta, V., Dalíková, M., Čapková Frydrychová, R., Neven, L. G., Sahara, K., & Marec, F. (2013). Neo-Sex Chromosomes and Adaptive Potential in Tortricid Pests. Proceedings of the National Academy of Sciences, 110(17), 6931–6936. 10.1073/pnas.1220372110

van Nieukerken, E. J., Kaila, L., Kitching, I. J., Kristensen, N. P., Lees, D. C., Minet, J., Mitter, C., Mutanen, M., Regier, J. C., Simonsen, T. J., Wahlberg, N., Yen, S. -H., Zahiri, R., Adamski, D., Baixeras, J., Bartsch, D., Bengtsson, B. Å., Brown, J. W., Bucheli, S. B., et al. (2011). Order Lepidoptera Linnaeus, 1758. In Z. -Q. Zhang (Ed.), Animal biodiversity: An outline of higher-level classification and survey of taxonomic richness (pp. 212–221). Magnolia Press, Auckland.

Ohno, S. (1966). Sex Chromosomes and Sex-linked Genes. Springer Berlin, Heidelberg.

Pazhenkova, E. A., & Lukhtanov, V. A. (2023). Chromosomal conservatism vs chromosomal megaevolution: enigma of karyotypic evolution in Lepidoptera. Chromosome Research, 31(2), 16. 10.1007/s10577-023-09725-9

Pennell, M. W., Kirkpatrick, M., Otto, S. P., Vamosi, J. C., Peichel, C. L., Valenzuela, N., & Kitano, J. (2015). Y Fuse? Sex Chromosome Fusions in Fishes and Reptiles. PLOS Genetics, 11(5), e1005237. 10.1371/journal.pgen.1005237

Pflu, J. M., Holmes, V. R., Burrus, C., Johnston, J. S., & Maddison, D. R. (2020). Measuring Genome Sizes Using Read-Depth, k-mers, and Flow Cytometry: Methodological Comparisons in Beetles (Coleoptera). G3 (Bethesda), 10(9), 3047-3060. 10.1534/g3.120.401028

de Prins, J., & Saitoh, K. (2003). Karyology and Sex Determination. In N. P. Kristensen (Ed.) Handbook of Zoology. Lepidoptera, Moths and Butterflies (pp. 449–468). De Gruyter.

Quinlan, A. R., & Hall, I. M. (2010). BEDTools: a flexible suite of utilities for comparing genomic features. Bioinformatics, 26(6), 841–842. 10.1093/bioinformatics/btq033

Robinson, R. (1971). Lepidoptera Genetics. Pergamon.

Sahara, K., Marec, F., & Traut, W. (1999). TTAGG Telomeric Repeats in Chromosomes of Some Insects and Other Arthropods. Chromosome Research, 7(6), 449–460. 10.1023/a:1009297729547

Schmitt, T., & Seitz, A. (2001). Allozyme Variation in *Polyommatus Coridon* (Lepidoptera: Lycaenidae): Identification of Ice-Age Refugia and Reconstruction of Post-Glacial Expansion. Journal of Biogeography, 28(9), 1129–1136. 10.1046/j.1365-2699.2001.00621.x

Šíchová, J., Nguyen, P., Dalíková, M., & Marec, F. (2013). Chromosomal Evolution in Tortricid Moths: Conserved Karyotypes with Diverged Features. PLoS ONE, 8(5), e64520. 10.1371/journal.pone.0064520

Speel, E. J., Hopman, A. H., & Komminoth, P. (2000). Signal amplification for DNA and mRNA. In I. A. Darby (Ed.), In Situ Hybridization Protocols (pp. 195–216). Humana Press.

Talavera, G., Lukhtanov, V. A., Pierce, N. E., & Vila, R. (2013a). Establishing Criteria for Higher-Level Classification Using Molecular Data: The Systematics of *Polyommatus* Blue Butterflies (Lepidoptera, Lycaenidae). Cladistics, 29(2), 166–192. 10.1111/j.1096-0031.2012.00421.x

Talavera, G., Lukhtanov, V. A., Rieppel, L., Pierce, N. E., & Vila, R. (2013b). In the Shadow of Phylogenetic Uncertainty: The Recent Diversification of *Lysandra* Butterflies through Chromosomal Change. Molecular Phylogenetics and Evolution, 69(3), 469–478. 10.1016/j.ympev.2013.08.004

Tomaszkiewicz, M., Medvedev, P., & Makova, K. D. (2017). Y and W Chromosome Assemblies: Approaches and Discoveries. Trends in Genetics, 33(4), 266–282. 10.1016/j.tig.2017.01.008

Traut, W. (1976). Pachytene Mapping in the Female Silkworm, *Bombyx Mori L*. (Lepidoptera). Chromosoma, 58(3), 275–284. 10.1007/BF00292094

Traut, W., Schubert, V., Daliková, M., Marec, F., & Sahara, K. (2019). Activity and inactivity of moth sex chromosomes in somatic and meiotic cells. Chromosoma, 128(4), 533–545. 10.1007/s00412-019-00722-8

Traut, W., Sahara, K., & Ffrench-Constant, R. H. (2023). Lepidopteran Synteny Units Reveal Deep Chromosomal Conservation in Butterflies and Moths. G3 (Bethesda), 13(8), jkad134. 10.1093/g3journal/jkad134

Van’t Hof, A. E., Marec, F., Saccheri, I. J., Brakefield, P. M., & Zwaan, B. J. (2008). Cytogenetic Characterization and AFLP-Based Genetic Linkage Mapping for the Butterfly *Bicyclus anynana*, Covering All 28 Karyotyped Chromosomes. PLoS One, *3*(12), e3882. 10.1371/journal.pone.0003882

Van’t Hof, E. A., Nguyen, P., Dalíková, M., Edmonds, N., Marec, F., & Saccheri, I. J. (2013). Linkage Map of the Peppered Moth, *Biston Betularia* (Lepidoptera, Geometridae): A Model of Industrial Melanism. Heredity (Edinb), 110(3), 283–295. 10.1038/hdy.2012.84

Vershinina, A. O., Anokhin, B. A., & Lukhtanov, V. A. (2015). Ribosomal DNA Clusters and Telomeric (TTAGG)n Repeats in Blue Butterflies (Lepidoptera, Lycaenidae) with Low and High Chromosome Numbers. Comparative Cytogenetics, 9(2), 161–171. 10.3897/CompCytogen.v9i2.4715

Vershinina, A. O., & Lukhtanov, V. A. (2017). Evolutionary Mechanisms of Runaway Chromosome Number Change in *Agrodiaetus* Butterflies. Scientific Reports, 7(1), 8199. 10.1038/s41598-017-08525-6

Vila, R., Lohse, K., Hayward, A., Laetsch, D. R., & Wright, C. (2023). The genome sequence of the Chalkhill Blue, *Lysandra coridon* (Poda, 1761). Wellcome Open Research, *12*(8), 162. 10.12688/wellcomeopenres.19253.1

de Vos, J. M., Augustijnen, H., Bätscher, L., & Lucek, K. (2020). Speciation through Chromosomal Fusion and Fission in Lepidoptera: Chromosomal Fusion & Fission. Philosophical Transactions of the Royal Society B: Biological Sciences, 375(1806), 20190539. 10.1098/rstb.2019.0539

White, M. J. D. (1946). The evidence against polyploidy in sexually reproducing animals. The American Naturalist, 80, 610–619.

Wiemers, M. (2003). Chromosome Differentiation and the Radiation of the Butterfly Subgenus *Agrodiaetus* (Lepidoptera: Lycaenidae: *Polyommatus*) - a Molecular Phylogenetic Approach [Dissertation]. Rheinische Friedrich-Wilhelms-Universität Bonn.

Wiemers, M., Stradomsky, B. V., & Vodolazhsky, D. I. (2010). A molecular phylogeny of *Polyommatus* s. str. and *Plebicula* based on mitochondrial *COI* and nuclear *ITS2* sequences (Lepidoptera: Lycaenidae). European Journal of Entomology, 107(3), 325–336. 10.14411/eje.2010.041

Winnepenninckx, B., Backeljau, T., & De Wachter, R. (1993). Extraction of High Molecular Weight DNA from Molluscs. Trends in Genetics, 9(12), 407. 10.1016/0168-9525(93)90102-n

Wright, C. J., Stevens, L., Mackintosh, A., Lawniczak, M., & Blaxter, M. (2024). Comparative Genomics Reveals the Dynamics of Chromosome Evolution in Lepidoptera. Nature Ecology & Evolution, 8(4), 777–790. 10.1038/s41559-024-02329-4

Yoshido, A., Marec, F., & Sahara, K. (2005). Resolution of Sex Chromosome Constitution by Genomic in Situ Hybridization and Fluorescence in Situ Hybridization with (TTAGG)n Telomeric Probe in Some Species of Lepidoptera. Chromosoma, 114(3), 193–202. 10.1007/s00412-005-0013-9

Yoshido, A., Marec, F., & Sahara, K. (2016). The fate of W chromosomes in hybrids between wild silkmoths, *Samia cynthia* ssp.: no role in sex determination and reproduction. Heredity (Edinb), *116*(5), 424-433. 10.1038/hdy.2015.110

Yoshido, A., Šíchová, J., Kristýna Pospíšilová, K., Nguyen, P., Voleníková, A., Šafář, J., Provazník, J., Vila, R., & Marec, F. (2020). Evolution of Multiple Sex Chromosomes Associated with Dynamic Genome Reshuffling in *Leptidea* Wood White Butterflies. Heredity (Edinb), 123(5), 138–154. 10.1038/s41437-020-0325-9

