## Supplementary material for "Polyommatine blue butterflies reveal unexpected integrity of the W sex chromosome amid extensive chromosome fragmentation"

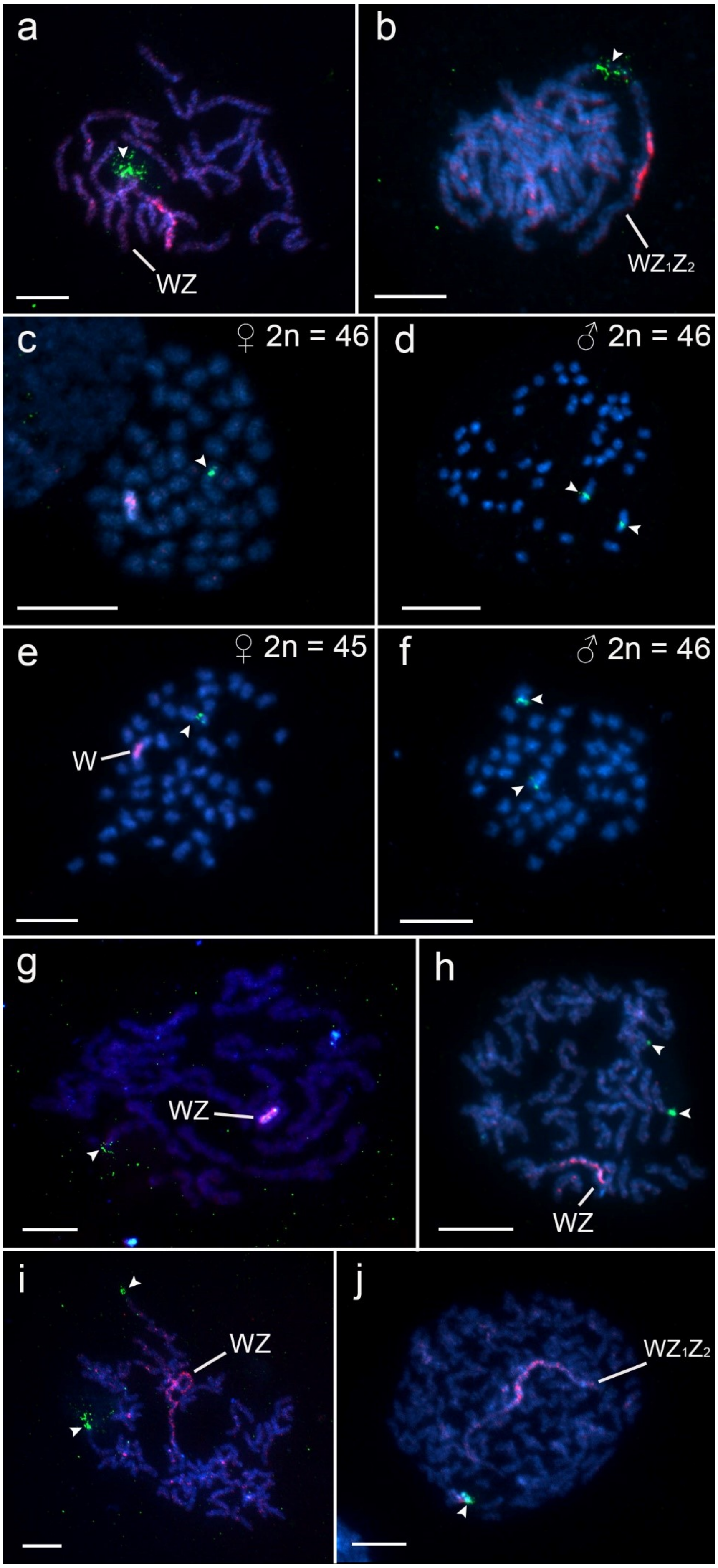
a
WZ
b
$WZ_1Z_2$
c
♀ 2n = 46
d
♂ 2n = 46
e
♀ 2n = 45
W
f
♂ 2n = 46
g
WZ
h
WZ
i
WZ
j
$WZ_1Z_2$

**Figure S1**. Genomic *in situ* hybridization and fluorescence *in situ* hybridization with 18S rDNA probe in pachytene (**a–b**, **g–j**) and mitotic (**c–f**) nuclei. Chromosomes were stained by DAPI (blue), the female gDNA was labelled with Cy3-dUTP or fluorescein dUTP (red) and 18S rDNA probe was labelled with biotin-16-dUTP detected by streptavidine conjugated with Cy3 (green). Arrowheads mark the rDNA loci. (**a**, **b**) Female pachytene nuclei of two karyotype races of *P. icarus* with 2n = 46 and 2n = 45, respectively. In both cases 18S rDNA probe hybridized to an interstitial site pairing with the euchromatic part of the chromosome W. (**c**) Female mitotic nucleus of *P. icarus* with 2n = 46. The rDNA locus was detected on the second largest chromosome, the Z chromosome, but not on the W. (**d**) Z-linkage of the rDNA locus is corroborated by two 18S rDNA signals in the largest chromosome pair in mitotic nucleus of a male from the same family as the female in (c). (**e**) Female mitotic nucleus of *P. icarus* with 2n = 45. (**f**) Mitotic nucleus of *P. icarus* male from the same family as the female in (e). (**g**) Female pachytene nuclus of *P. escheri* with a single terminal autosomal rDNA locus. (**h**) Female pachytene nucleus of *L. bellargus* with two terminal autosomal rDNA loci. (**i**) Female pachytene nucleus of *L. coridon* with the terminal 18S rDNA loci located on two autosomal bivalents. (**j**) Female pachytene nucleus of *P. dorylas* with a single terminal 18S rDNA cluster located on one autosomal bivalent. Scale bar 10 μm.

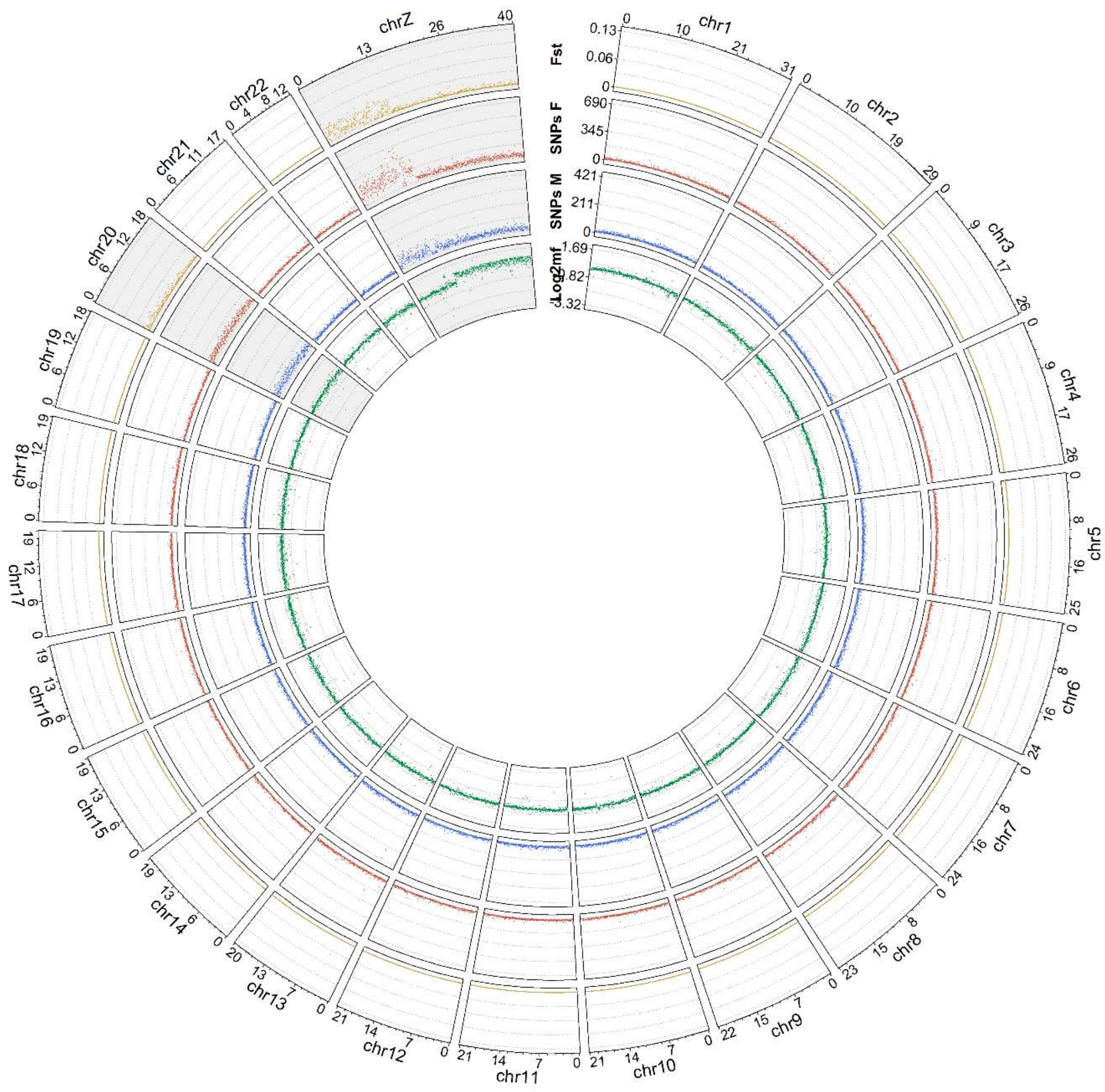

**Figure S2**: Identification of sex-linked regions in the *P. icarus* race with $WZ_1Z_2$ sex chromosomes. Circo plot shows results of Coverage (green) and Pool-seq analyses (male- and female-specific SNPs and their relative divergence - $F_{st}$ are in blue, red and yellow, respectively). The Z chromosomes are shaded in grey.
